# Efficient inference of evolutionary and progressive dynamics on hypercubic transition graphs

**DOI:** 10.1101/2022.05.09.491130

**Authors:** Marcus T. Moen, Iain G. Johnston

## Abstract

The progression of cancer and other diseases, the evolution of organismal features in biology, and a wide range of broader questions can often be viewed as the sequential stochastic acquisition of binary traits (for example, genetic changes, symptoms, or characters). Using potentially noisy or incomplete data to learn the sequences by which such traits are acquired is a problem of general interest. The problem is complicated for large numbers of traits which may, individually or synergistically, influence the probability of further acquisitions both positively and negatively. Hypercubic inference approaches, based on hidden Markov models on a hypercubic transition network, address these complications, but previous Bayesian instances can consume substantial time for converged results, limiting their practical use. Here we introduce HyperHMM, an adapted Baum-Welch (expectation maximisation) algorithm for hypercubic inference with resampling to quantify uncertainty, and show that it allows orders-of-magnitude faster inference while making few practical sacrifices compared to existing approaches. We apply this approach to synthetic and biological datasets and discuss its more general application in learning evolutionary and progressive pathways.

## Introduction

Many questions in biology, medicine, and beyond concern the dynamics by which a set of traits or ‘characters’ is acquired over time. These traits could be, for example, physiological characters in evolutionary biology, mutations in cancer progression, symptoms in progressive disease, task completions in a workflow, and more. Efficient ways of learning about these dynamics from available data – which may be single-time snapshots, without longitudinal tracking of individuals – can be challenging to implement.

Specific fields of study have given rise to different approaches to this question. The field of cancer evolution [1] has developed methods focussing on the construction of mutation graphs describing the ordering and dependence of mutational changes. Several of these approaches use a Bayesian networks picture [2, 3, 4, 5, 6, 7, 8], which may describe dependencies between mutations as deterministic and one-way (i.e. detecting when X is required for Y, but not when Z has the effect of lowering the probability of Y) – as described, and relaxed, in recent work considering a different type of model called mutual hazard networks [9]. Other approaches for cancer progression have been developed that use alternative methods based, for example on the analysis of permutations [10, 11], and Markov modelling [12]; meta-studies have compared the performance of several of these approaches [13]. In the (not disconnected) field of evolutionary biology, several approaches have been developed for describing and predicting the appearance of traits (typically called ‘characters’ in an evolutionary setting) on phylogenies. In an evolutionary setting, the combined problem of inferring character evolution on a phylogeny and the phylogeny structure itself is often considered [14, 15]. Bayesian approaches for phylogenetic reconstruction [16, 17] are often combined with Markov models for character dynamics [14], describing the different states of a character or characters and allowing stochastic transitions between those states with some rates that are model parameters to be estimated. Recent developments including generalising the influences between evolving characters to include dependence and conditionality [18], employing flexible hidden Markov models to describe character dynamics [19], and simulation-free approaches allowing computationally tractable treatment of problems involving ensembles of possible trees [20]. The links between the cancer and evolutionary fields have been explicitly explored by several studies picturing cancer progression as an evolutionary process [21, 22].

In parallel, several studies have considered a particular class of applied problems which we will call ‘hypercubic inference’, applied in cancer progression [22, 9], evolutionary biology [22, 23, 24], and progression of other diseases including severe malaria [25]. In terms of the systems involved, this picture involves evolution progressing via the sequential, irreversible, stochastic acquisition of discrete traits (also referred to as monotonic accumulation [2]). Rather than focussing on individual traits/characters as the elements of the system, these approaches consider every possible state of a system involving *L* traits – thus explicitly considering every combination of trait values, and thus accounting for completely general influences of any subset of current trait values and the stochastic acquisition of another. There are therefore no assumptions of deterministic or one-way relationships [22, 9]. The transition graph linking possible states is then a directed hypercubic graph, and edge weights (model parameters to be estimated) can be used to control the probabilities of different dynamic pathways (Fig. 1). We are concerned with the inverse problem of learning the structure of, and variability in, pathways on thus hypercube from observed samples of the evolving system. The set of observations used to parameterise the hypercubic model may, in different scientific contexts, be cross-sectional, longitudinal, or phylogenetically coupled [22].

**Figure 1:**
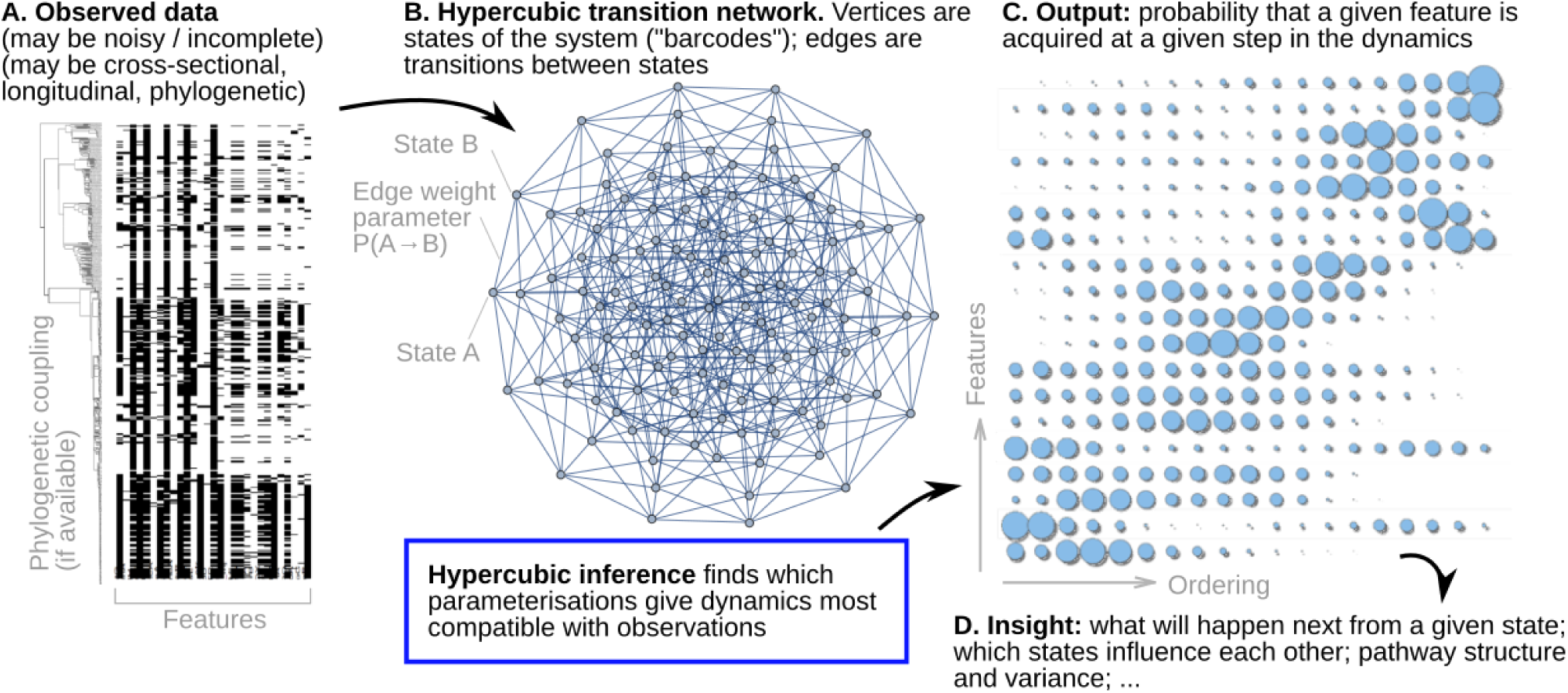
Overview of hypercubic inference. (A) Observed data in the form of presence/absence ‘barcodes’ for each observation, which may be incomplete or noisy, and may be independent (cross-sectional), longitudinal, or phylogenetically coupled. (B) The hypercubic transition network model for dynamics, where a system proceeds via a series of transitions from one vertex to another. Each vertex is a different ‘barcode’ state, edges give transition probabilities between states. Hypercubic inference learns these transition probabilities from data, finding the parameterisation most compatible with a set of emitted observations. (C) The learned parameterisation can be interpreted in several ways – as a probability map of which feature is likely acquired at which stage, explicit pathways through the hypercube space, relationships betwene feature orderings, and more. (D) Scientific insight follows from interpreting these results.

Hypercubic transition path sampling (HyperTraPS) [23, 22] and its precursor phenotypic landscape inference (PLI) [26] are Bayesian approaches, using likelihood estimates to build posterior distributions over the parameterisations of the hypercube. The representation of parameters is flexible, with a function used to construct each edge weight from some potentially lower-dimensional representation, including the proportional hazards picture also used in Ref. [9]. Regularisation approaches have been used to seek the parameter representation most compatible with given data, which itself informs on the generative relationships between features [22]. HyperTraPS has been used to infer the evolution of efficient photosynthesis in plants [26], gene loss in mitochondria [23], as well as the progression of severe malaria [25], the emergence of tool use in animals [24], and the participation of students in digital learning [27].

While general, this approach, which relies both on Markov chain Monte Carlo and a sampling approach to estimate likelihoods, is computationally expensive and approximate. The inclusion of prior information is natural and arguably important for low sample sizes, to avoid overfitting a small sample. However, for larger samples, we may expect lower influence of priors on the posterior. We may also wish to avoid the Bayesian paradigm outright. We can thus naturally ask if a computationally cheaper approach can provide an output akin to a maximum likelihood estimate for hypercube parameters. Here, we will develop and apply HyperHMM, an alternative approach for inference of dynamic pathways on directed hypercubes.

## Results

### A hypercubic Baum-Welch algorithm for efficient inference of paths on a transition hypercube

Under a hypercubic transition graph model, every possible state of the system is represented as a binary string of length *L*, where 0 and 1 at the *i*th position correspond, respectively, to absence or presence of the *i*th trait (Fig. 1). Traits are acquired stochastically and irreversibly, meaning once a trait has been acquired it can not be lost [22]. A hypercubic transition graph is then constructed, with each node corresponding to a state, and a weighted edge from node *a* to node *b* if *b* differs from *a* by the acquisition of exactly one trait. We usually picture a given instance of the system (for example, a patient with a progressive disease) moving over the hypercube from the binary string of all 0s towards (but not necessarily reaching) the binary string of all 1s, probabilistically following outgoing edges from a given state according to their relative weights. The goal of hypercubic inference is to learn the set(s) of edge weights that are most compatible with the observed dynamics of a given system. To this end, we consider a hidden Markov model (HMM) likelihood based on emissions from processes on this transition graph [28]. In the simplest case, an emission corresponds simply to the state at the current node, but an HMM approach also allows us to account for noisy and incomplete emissions.

Hypercubic transition path sampling (HyperTraPS) is an algorithm estimating the likelihood of a given observed transition from some state *a* to some state *b* (not necessarily only one trait apart). Given the large number of paths that can generally exist between two nodes, HyperTraPS uses biased random walkers to estimate this likelihood, which is then embedded in a Bayesian framework using Markov chain Monte Carlo (MCMC) for parameter estimation. The fact that this likelihood is approximate raises issues of the MCMC convergence which require corresponding algorithmic complexity to address [22, 29] and the Bayesian nature of the parameter search require substantial computer time.

Here, we propose HyperHMM, an alternative using (a) an adaptation of the well-known Baum-Welch algorithm [30, 28] for the hypercubic transition graph to estimate parameters without requiring an approximate likelihood, and (b) a frequentist approach using resampling rather than the fully Bayesian approach for uncertainty quantification. Both (a) and (b) allow substantial computation gains over the usual implementation of HyperTraPS, reducing runtimes from hours to seconds (see below). The Baum-Welch algorithm is an expectation-maximisation algorithm that learns maximum likelihood transition probabilities in an HMM from a given set of observations, and provides the maximum-likelihood counterpart to the Bayesian approach previously used with HyperTraPS [23, 22]. The core hypercubic Baum-Welch algorithm is given in Algorithm 1, illustrated in Fig. 1, and derived in the Methods and Supplementary Information.

#### Algorithm 1

The Baum-Welch algorithm for inference on hypercubic transition graphs. The form of the *P* (…) functions is given in the Methods section.

Input: *L* = The number of traits, and *O* = *O*_1_, …, *O*_*R*_ = all the independent observations

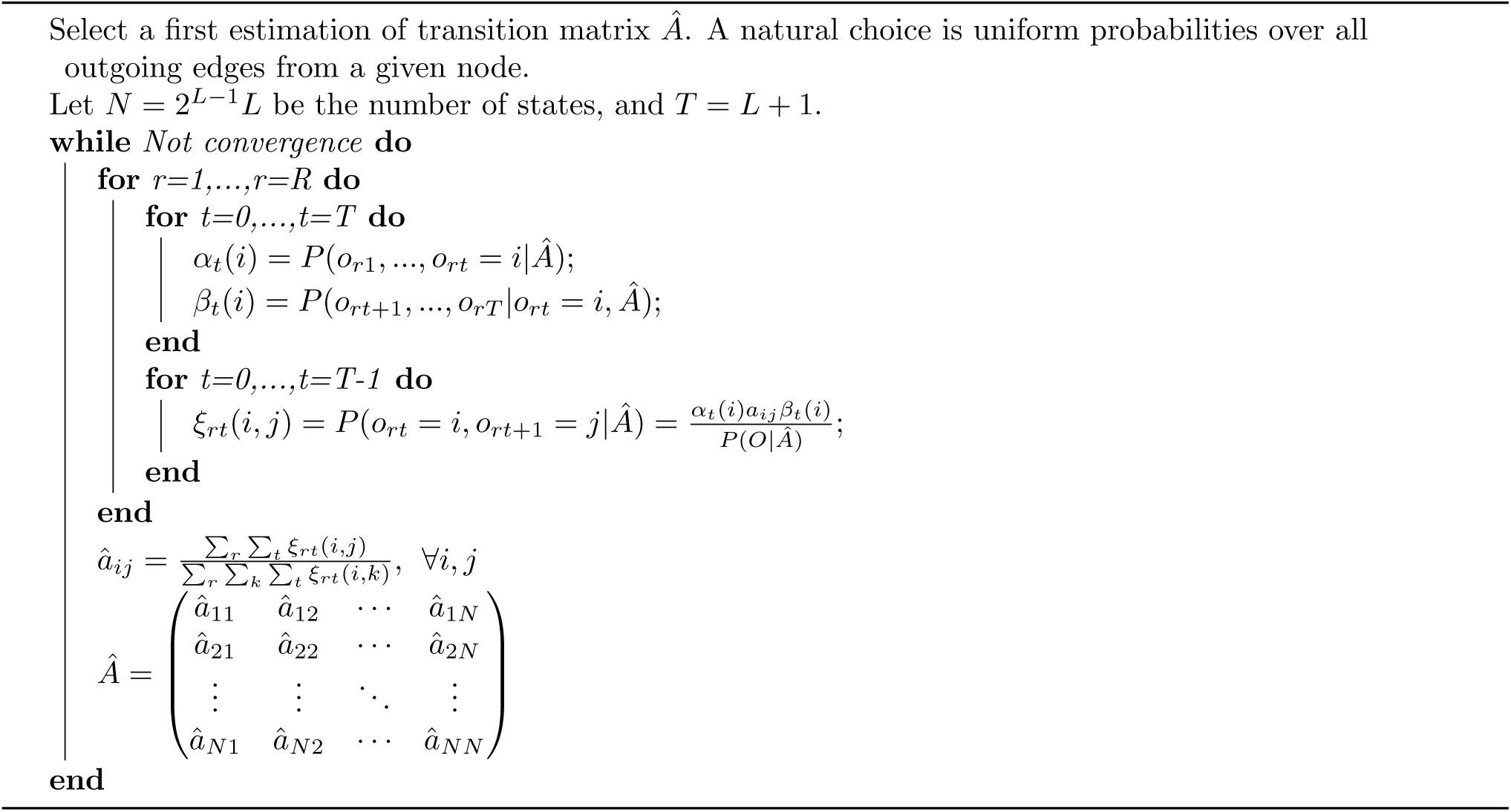

### Synthetic test cases – single and competing evolutionary pathways

We will start by looking at two synthetic cases involving simple preconstructed datasets, previously used to test HyperTraPS [22]. In the first case, we have constructed a dataset reflecting a single evolutionary pathway, where the acquisition of features proceeds from the first to the last indexed. We begin with this simple system with *L* = 5 traits, with corresponding ‘data’ consisting of the observations 10000, 11000, 11100, 11110, repeated *N* times to demonstrate the behaviour of inferred parameters with increasing dataset size.

Fig. 2A shows the inferred probabilities with which a feature is acquired in a given order, for different dataset size *N*, and for the HyperHMM and Bayesian HyperTraPS approaches. The single pathway structure is intuitively captured, with increasing certainty as the dataset size increases. The uncertainty derived from resampling with the HBW algorithm agrees well with the posteriors from HyperTraPS. HyperTraPS, as a Bayesian approach, is informed by its priors, which in this case are simply uniform acquisition probability over all options. For low *N* the influence of these priors on the posteriors is greater, and corresponding the uncertainty quantified from HBW resampling is higher.

**Figure 2:**
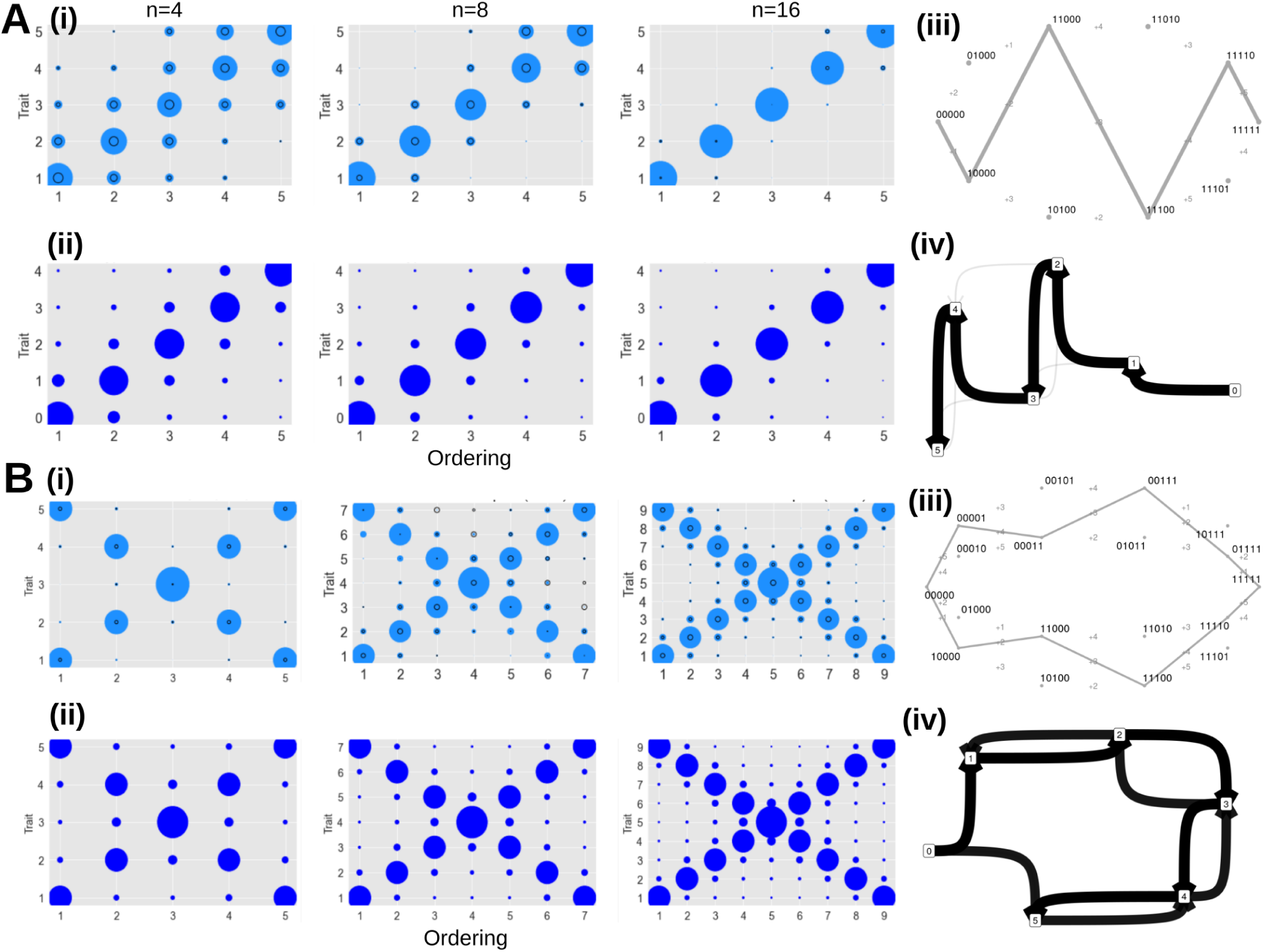
Inferred dynamics for synthetic test datasets. (A) Single pathway with varying sample size, (B) double pathway with varying feature count examples from the text. (i) Summary output of HyperHMM algorithm for different sample size on each dataset; (ii) summary HyperTraPS posteriors. Blue bubbles show the probability of getting trait *y* at time *x*; black circles in the HyperHMM plots shows the standard deviation after 100 bootstraps. (iii) Visualisation of inferred paths on the hypercubic transition network. Individual edge labels describe which feature is changed at each transition; edge weights correspond to the probability of a given transition. (iv) Probabilistic feature graphs for orderings of features changes. An edge from *a* to *b* corresponds to acquisition of *b* directly following acquisition of *a* in inferred dynamics; 0 gives the initial state with no features. Edge weights correspond to the frequency with which given transitions are observed in simulated dynamics.

To demonstrate the algorithm’s ability to capture distinct pathway structures, and negative interactions between traits, we next consider a case with two competing pathways. Here, the first trait to be acquired is equally likely to be the first or last indexed. If the first, evolution proceeds as previously, but if the last, it proceeds in the ‘opposite direction’, with traits being acquired last to first in indexing order. The first step on each pathway thus represses the other pathway. The HyperHMM probabilities and HyperTraPS posteriors are shown in Fig. 2B, this time varying the number of traits *L* in the dataset. Both algorithms capture the double pathway structure of the data as before, and the structure of uncertainty in both cases is the same.

### Ovarian cancer data

Following the verification of the HyperHMM approach on synthetic datasets, we next turned to a medical dataset that, while dated, has been used to test several algorithms for evolutionary inference [22, 8, 4]. This dataset consists of snapshots of patterns of chromosomal aberrations in 87 ovarian cancer patients [31]. These are labelled by chromosome (1 *−* 23 and *X*), chromosomal arm (*p* or *q*) and variant type (addition + or loss *−*). HyperTraPS and others have used these data to test and benchmark inference approaches [22]. The questions include: do some aberrations occur systematically before others, does the presence of one aberration influence the acquisition probability of another, and how distinct or separate are the different pathways by which cancer can evolve in this dataset?

Fig. 3 shows the inferred orderings from the HyperHMM approach and from HyperTraPS. The same features are clear in both cases, with sampled hypercubic trajectories and probabilistic feature graph (Fig. 3 C-D) displaying notable similarities with the HyperTraPS output found in Ref. [22] – including strong weighting to the 8*q*+ → 3*q*+ → 1*q*+ → 4*q−* sequence, then a more diverse range of potential dynamics thereafter. The supported trajectories here clearly underline the ability of HyperHMM to characterise competing pathways (and negative influence of one trait on the acquisition probability of another). As discussed in Ref. [22], this leads to increased flexibility in pathway identification compared to other approaches: for example, the HyperHMM output supports acquisition of 4*q−* prior to 5*q−*, observed in 12% of samples in the data but not identified by inference approaches based on Bayesian networks [5, 7]. More generally, hypercubic inference approaches allow a relaxation of the graph of trait relationships away from the tree structure enforced by many approaches to allow bidirectional interactions; and they also capture more detailed information about transitions between individual states, allowing general representations of trait dependencies.

**Figure 3:**
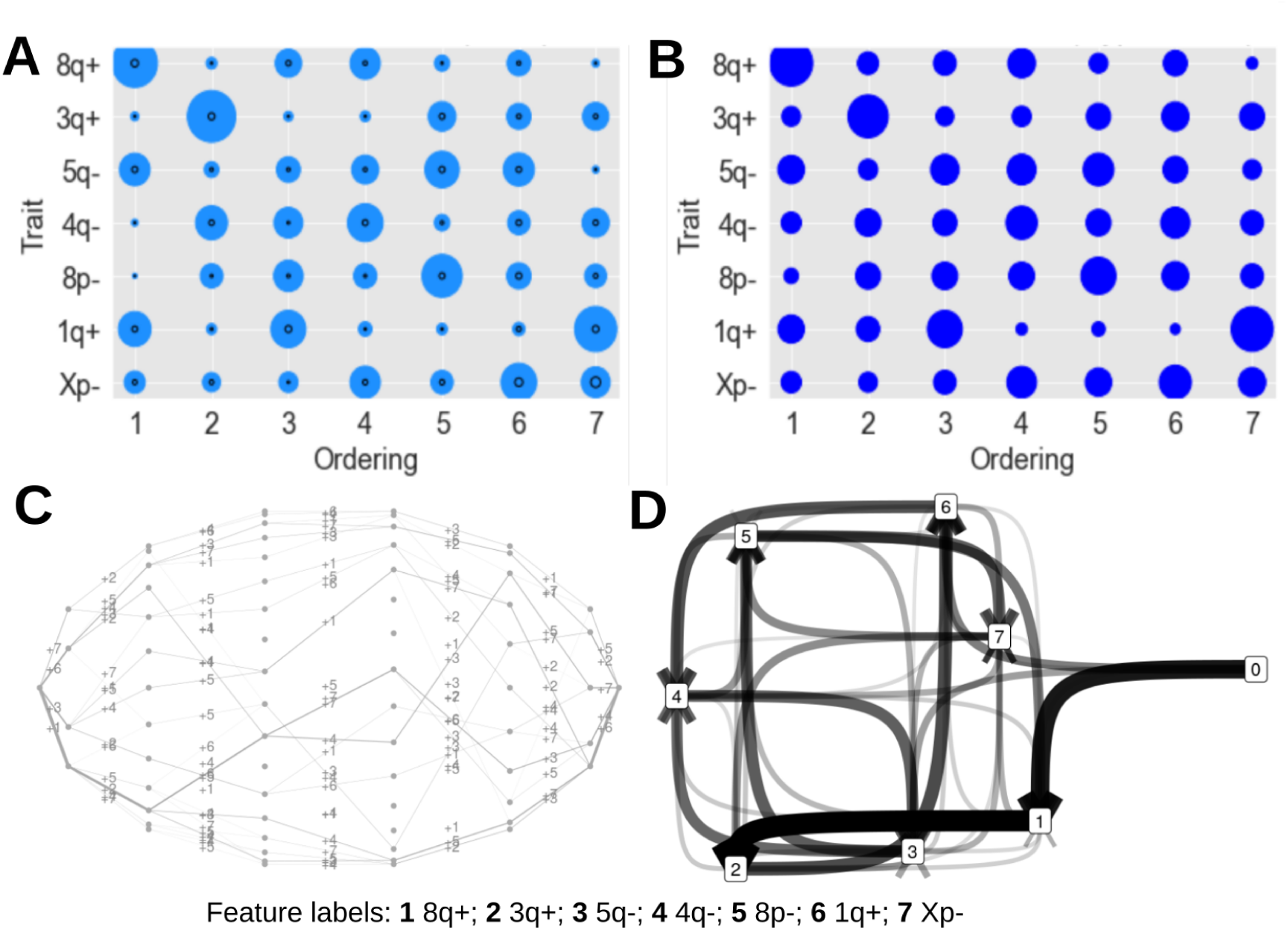
Inferred dynamics for ovarian cancer progression. (A) Summary results using the HyperHMM algorithm and (B) using HyperTraPS on the ovarian cancer dataset. Bubbles show the probability of getting trait *y* at time *x*; black circles in the HyperHMM plots show the standard deviation after 100 bootstraps. (C) Visualisation of inferred paths on the hypercubic transition network. Individual edge labels describe which feature is changed at each transition; edge weights correspond to the probability of a given transition. (D) Probabilistic feature graphs for orderings of features changes. An edge from *a* to *b* corresponds to acquisition of *b* directly following acquisition of *a* in inferred dynamics; 0 gives the initial state with no features. Edge weights correspond to the frequency with which given transitions are observed in simulated dynamics.

Comparing Figs. 3 A-B shows that the HyperTraPS output is more influenced by the uniform priors applied in that analysis. The distribution of acquisition probabilities across features and orderings is more uniform compared to the HyperHMM output, which distributes probability mass more heterogeneously. There is likely another contributing factor to this difference, reflecting the face that HyperHMM is capturing a single maximum likelihood parameterization (per bootstrap resample) and that the Bayesian approach’s wider exploration of parameter space identifies some parameterisations that lead to greater uncertainty (this is more likely in the case of complex real data than in the synthetic cases above, where a single optimal parameterization can capture all the dynamics). In this case, commonly found in frequentist-Bayesian comparisons, the Bayesian picture will give a more accurate reflection of the true uncertainty under a particular set of priors, but the HyperHMM solution is much more cheaply available and readily captures the essential dynamic features.

### Multi-drug resistance in tuberculosis

As a final test of the approach using biomedically pertinent data, we next turned to an evolutionary question – how multidrug resistance evolves in tuberculosis (TB). This societal problem involves the tuberculosis bacterium acquiring resistance to the antimicrobial drugs used to treat it. We again use a dataset that has previously been used to test HyperTraPS [22]: specifically, the Casali *et al*. study of Ref [32]. Here, drug resistance profiles of different but related strains of TB were experimentally characterised, producing barcodes of resistance/susceptibility that are connected by an estimated phylogeny. Following Ref. [22], we estimate the barcodes of ancestral states using maximum parsimony and hence extract the set of transitions that have occurred throughout the whole lineage. This transition set is the input for our inference algorithms.

Once more, the structure and variability of multidrug resistance pathways is readily revealed by the HyperHMM algorithm. Uncertainty quantification recapitulates the fully Bayesian approach using HyperTraPS (which did not yield fully converged posteriors for this case). Following findings in Ref. [22], a strong weighting to the initial steps INH → RIF → PZA is apparent, after which the potential pathways diversify around a central ‘core’ pathway. Interestingly, this suggests that acquisition of multidrug resistance in each lineage in these samples follows one of a related set of evolutionary pathways, with some variability around later steps but the same qualitative patterning (there are not multiple, distinct, competing pathways in Fig. 4C to the extent seen in Fig. 2B). This raises the possibility of predicting the next evolutionary step for a strain with a given drug resistance profile, with potential applications to optimal treatment design (akin to Ref. [25]).

**Figure 4:**
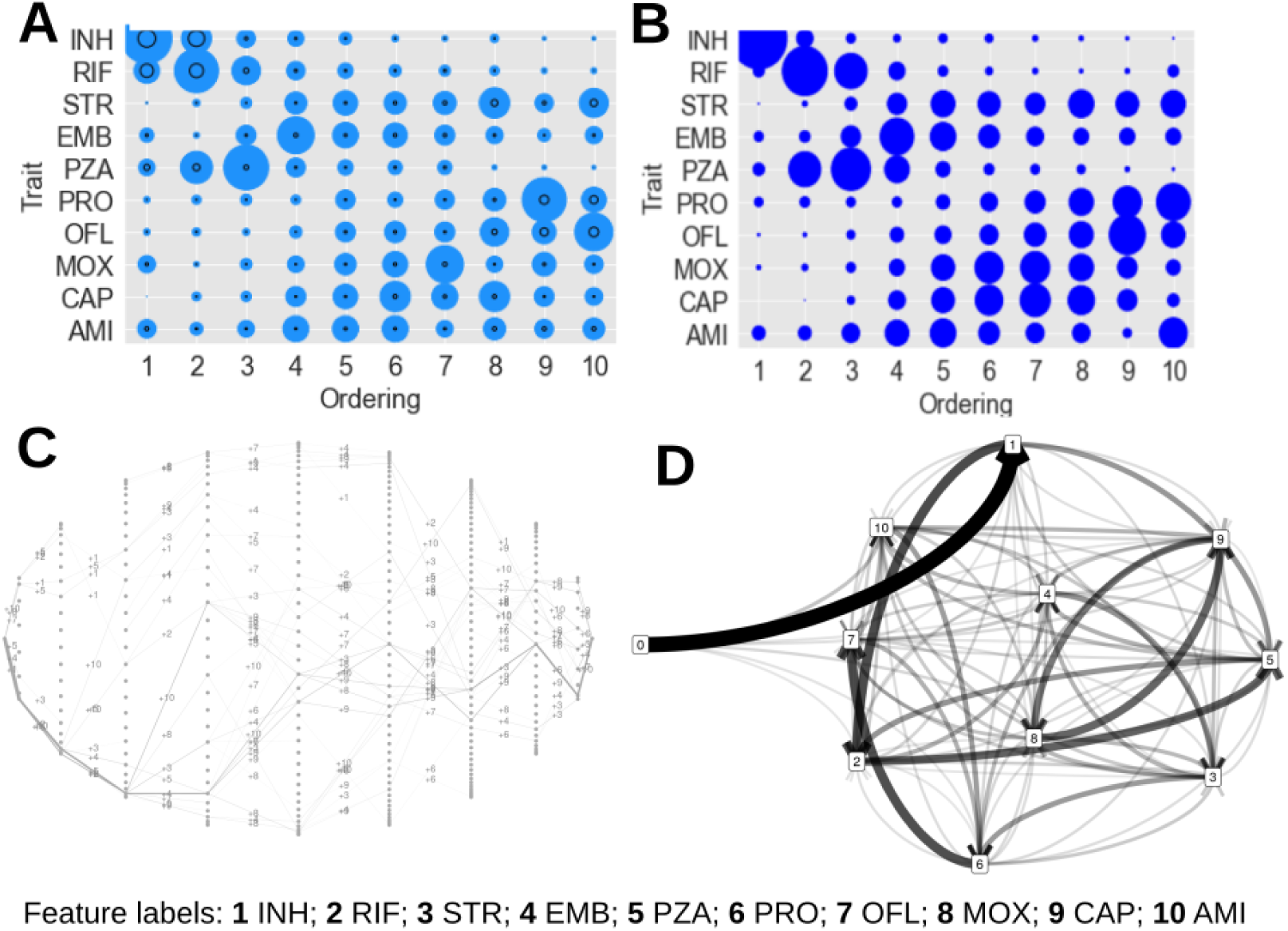
Inferred dynamics for multidrug resistance evolution in tuberculosis. (A) Summary results using the HyperHMM algorithm and (B) using HyperTraPS on the tuberculosis drug resistance dataset. Each three-letter code is a particular drug. Bubbles show the probability of getting trait *y* at time *x*; black circles in the HyperHMM plots show the standard deviation after 100 bootstraps. (C) Visualisation of inferred paths on the hypercubic transition network. Individual edge labels describe which feature is changed at each transition; edge weights correspond to the probability of a given transition. (D) Probabilistic feature graphs for orderings of features changes. An edge from *a* to *b* corresponds to acquisition of *b* directly following acquisition of *a* in inferred dynamics; 0 gives the initial state with no features. Edge weights correspond to the frequency with which given transitions are observed in simulated dynamics.

### Benchmarking performance and comparison to Bayesian approach

Throughout the above investigations, we observed that the hypercubic Baum-Welch algorithm yielded results much more quickly than the HyperTraPS approach – intuitively, as a maximum-likelihood approach will typically involve less computation than Bayesian inference attempting to summarise the full parameter space. To quantify the difference in speeds, we considered simple synthetic test cases of the single- and double-pathway systems introduced above with *L* = 5, 7, 9. In each case we used a synthetic dataset providing exactly two samples of every non-trivial transition on the corresponding hypercube. For the HyperHMM approach, we recorded how long it took for the inferred parameter set to converge to a 0.001 level, with 100 bootstraps to quantify uncertainty. For HyperTraPS, we recorded how long it took for the inferred ordering posteriors to converge to a 0.001 level. We did not focus on the posteriors on individual transition parameters, because many of these are unconstrained by a given dataset and (in the absence of a sparsity prior) take a long time to recapture the prior, for negligible contribution to the inferred dynamics. In a sense this criterion allows HyperTraPS a looser definition of convergence; however, even given this laxity, the speedup from the HyperHMM approach is striking (Table 1). It typically converges two to three orders of magnitude more quickly than the Bayesian implementation of HyperTraPS, without requiring preliminary or parallel tuning of the MCMC parameters.

**Table 1:** Convergence times for different hypercubic inference approaches. Timing results for various datasets using the HyperHMM algorithm and HyperTraPS. All times are in seconds; convergence criteria *E* are the maximum change to any probability in the model allowed between instances (HBW iterations; HyperTraPS sample blocks). The HyperHMM column shows the time it took to run 100 bootstraps with a convergence criterion of 0.001.

| Dataset | Convergence time / s |  |  |
| --- | --- | --- | --- |
| | Hypercubic Baum-Welch ($\epsilon = 0.001$) | HyperTraPS ($\epsilon = 0.001$) | HyperTraPS ($\epsilon = 0.01$) |
| $L = 5$ single | 0.0960 | 961 | 168 |
| $L = 7$ single | 0.768 | 3380 | 950 |
| $L = 9$ single | 3.70 | 8900 | 1308 |
| $L = 5$ double | 0.239 | 1181 | 305 |
| $L = 7$ double | 1.55 | 5914 | 1081 |
| $L = 9$ double | 6.06 | 16978 | 2521 |
| Ovarian | 19.3 | $> 2 \times 10^5$ | $> 2 \times 10^5$ |
| TB drug | 3480 | $\gg 2 \times 10^5$ | $> 2 \times 10^5$ |

### Prior information and model selection

One sacrifice in going from the fully Bayesian HyperTraPS is a reduced ability to include prior information in the inference process [22]. However, we can make some progress by encoding prior information in the structure of the hypercubic network that underlies our model. Say we know that trait *a* always appears before trait *b*. A simple way of including this prior information is by removing all edges in the hypercubic transition network that correspond to an acquisition of *b* from a state without trait *a*. The possibility of such a transition is thus removed from the inference process, and the HBW algorithm will identify the maximum likelihood parameterisation of the remaining edges given the observed data.

This approach of removing edges also allows a degree of model selection to be applied. We can consider different models for a given system, involving hypercubic networks with different edge sets removed. The HBW algorithm will identify a maximum likelihood parameterisation for each (or, if a chosen model is incompatible with observations, fail to identify any parameterisation and give a zero likelihood). The number of free parameters associated with a transition graph *G* is the number of outgoing edges *d*_*out*_(*s*) from each non-terminal node *s* minus one, summed over all nodes:

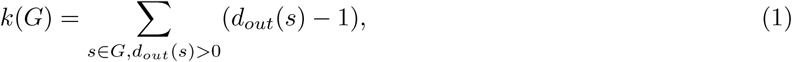

*k*(*G*) for the full hypercube can readily be computed as 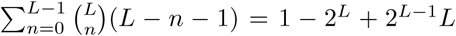. Given this parameter count and the associated likelihood, we can then use model selection approaches like the Akaike and Bayesian Information Criteria to choose parsimonious hypercube structures that are compatible with observations. As a simple example, consider the single and double pathway models above. We can consider three different model structures: (a) the full hypercube (as in all previous sections); (b) the hypercube with all edges removed except those on the single pathway 000… → 100… → 110… → …; (c) the hypercube with all edges removed except those on the two pathways 000… → …001 → …011→ … Then consider finding the maximum likelihood parameterisations of each of these models with (i) *N* samples from the single pathway system and (ii) *N* samples from the double pathway system. The results, along with associated AIC values, are given in Table. 2, demonstrating that models that are unnecessarily complex (a) or insufficiently complex (b)(ii) are penalised in favour of those that better match the observational structure (b)(i), (c)(ii).

**Table 2:** Model selection for different *L* = 5 experiments. We use the Akaike Information Criterion *AIC* = 2*k −* 2 log ℒ, where ℒ is likelihood and *k* is number of parameters. Rows give different synthetic datasets; columns give different parameter structures for the hypercubic model. In each case the most parsimonious model capable of capturing the data is favoured.

| | (a) full hypercube $k = 49$ | (b) single pathway $k = 0$ | (c) double pathway $k = 1$ |
| --- | --- | --- | --- |
| (i) single pathway | $\mathcal{L} \simeq 0.98$; $AIC \simeq 98$ | $\mathcal{L} = 1$; $AIC = 0$ | $\mathcal{L} = 1$; $AIC = 2$ |
| (ii) double pathway | $\mathcal{L} \simeq 1.45 \times 10^{-5}$; $AIC \simeq 120$ | $\mathcal{L} = 0$; $AIC = \infty$ | $\mathcal{L} = 0.5^{16} = 1.52 \times 10^{-5}$; $AIC \simeq 24$ |

The principle of edge removal can also be used to regularise identified models, as demonstrated in Ref. [22]. Here, the full hypercube model is first parameterised, then those edges with the lowest weights are removed and a model selection criterion recalculated. The process continues to iteratively remove low-probability edges until an unacceptable likelihood penalty results. While not flawless (for example, the best solution might involve removing a combination of edges that is not encountered in the iterative one-at-a-time approach), this approach can be used to lower the complexity of learned models given a set of observations.

## Discussion

We have shown that a hypercubic Baum-Welch algorithm is an efficient alternative for inferring dynamic pathways on hypercubic transition graphs, and can be combined with resampling for uncertainty quantification in the HyperHMM approach. The considerable speedup afforded over the original HyperTraPS implementation increases the set of problems that can be addressed; at the same time, no case-specific choices about perturbation kernels and likelihood estimation parameters need to be made. This simplification and speedup does not challenge the approach’s ability to capture and quantify uncertainty, making it a highly competitive choice for inferring the structure of, and variability in, dynamic pathways.

The ability to incorporate prior data in a Bayesian sense is lacking in this approach. However, as the algorithm considers all transitions between all states, one possibility is to restrict some of the transition edge weights to be zero, as we demonstrate with simple examples above. Applied systematically, this would allow, for example, the enforcement of a particular trait being acquired before another, by setting to zero any edge that disobeys this pattern. Another feature of the Bayesian approach – integration over the complete parameter space – is necessarily lacking in the HyperHMM approach and may lead to differences in cases where distinct parameter sets support similar behaviours (a rugged likelihood surface). However, the good comparison between HyperHMM and HyperTraPS for several cases above with complex and distinct pathway structures suggests that this may not be a fatal challenge in real datasets.

We have demonstrated uncertainty quantification via bootstrap resampling. This can be a computationally demanding process, scaling with the number of bootstrap resamples. Several other options exist that could potentially be employed. As the HBW algorithm returns what is a maximum likelihood estimate for the transition graph weights, numerical or automatic differentiation [33] could be used to calculate likelihood derivatives and thus provide confidence intervals on these parameters in future applications.

One advantage of the likelihood estimation process here is that noisy and incomplete observations can very easily be included in the source data, via appropriate choices for emission probabilities in the algorithm. Accounting for noisy observations is achieved by assigning each state a nonzero probability of emitting a signal that differs from its signature. The choice of this probability can be informed by the particular system, and could involve, for example, a constant error probability ϵ per bit, so that the probability of emitting a signal that differs by *l* bits from the current state is ϵ^*l*^ (appropriately normalised). Incomplete observations can be modelled by assigning each state compatible with a signal an equal probability of emitting that signal (for example, 100 and 110 may emit 1?0 with equal probability). This emission probability does not itself influence the probability of those states arising in the dynamics, which is determined by the inferred parameterisation.

The hypercubic Baum Welch algorithm considers every transition edge on the hypercube. In the examples above, because of the dramatic increase in computational efficiency, we have not coarse-grained the associated parameter space in any way. However, faced with larger problems, the same strategies for dimensionality reduction that were employed with HyperTraPS [22] – and have been employed in similar hypercubic models could readily be employed here. This would involve a potentially simple function in Algorithm 1 mapping a reduced parameter set *θ*′, for example involving a proportional hazard mapping as in [22, 9] to transition edge weights, and its inverse being used in the parameter update step each iteration.

Comparisons with alternative approaches for inferring dynamics in a similar way to the hypercubic picture have been shown in Refs. [23] and [22]. Classes of approach for this problem broadly include simple logistic regression, approaches based on Bayesian networks, approaches for modelling and learning stochastic processes on phylogenies, and approaches harnessing topology and/or dimensionality reduction before performing inference [22]. Of particular note are the approaches of Hjelm *et al*. [12] where a Markov chain picture was used to construct networks describing longitudinal observations with pairwise trait interactions; and of Schill *et al*. [9], who apply a different set of inference approaches to a very similar hypercubic model setup. Previous comparisons have demonstrated that HyperTraPS typically has advantages of scaling and generality over several other approaches. Our aim here is to demonstrate that the hypercubic Baum-Welch algorithm matches HyperTraPS output – and hence retains these advantages – while allowing a considerable speedup that will render more large-scale problems computationally tractable.

## Methods

### Hypercubic Baum-Welch algorithm

The derivation and intuition behind the hypercubic Baum-Welch algorithm is given more fully in the Supplementary Information. Here we simply state the essential aspects. We are concerned with estimating the probabilities *a*_*ij*_ = *P* (*X*_*n*_ = *s*_*j*_|*X*_*n−*1_ = *s*_*i*_), for a stochastic process *X*_*t*_ on the hypercubic graph where each state *s* is a node and edge *s*_*i*_ → *s*_*j*_ exists iff *j* differs from *i* by exactly one feature acquisition (Fig. 1. The algorithm broadly considers the set of possible paths on the hypercube that could lead to observations being ‘emitted’ that are compatible with our observations (see Fig. S1 for examples), and seeks to find transition weights that maximise the probability of these paths.

The data we will use is a set of potentially sequential observations *O*, where *O*_*r*_ = *o*_*r*0_, …, *o*_*rT*_, is the *r*th sequence of observation, each labelled by an observation ordering 0, …, *T*. Any of these observations may be absent.

The key idea is to find probabilities, *a*_*ij*_, that maximize the likelihood of seeing all of our observations. This is done by first calculating the probabilities going forward and backwards in time and storing the values for each time step. The forward probability is the probability of seeing everything up to a given time *t* given our current estimate of the transition matrix, and the backward probability is the probability of seeing everything from a given time *t* until the end. These quantities are defined as:

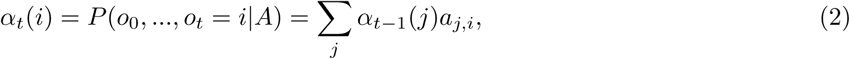

where *α*_0_(0^*L*^) = 1 and *α*_0_(*i*) = 0 for all other states *i*, and

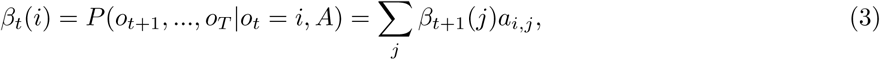

where *β*_*T*_ (*i*) = 1 for all states *i*.

Combining these probabilities will give us a ‘skeleton’ with all possible pathways given the observation (Fig. S1). Using this we can calculate the probability of being in any two states at any given time and update the transition matrix based on this information. These probabilities are called the *ξ*-probabilities and are defined as:

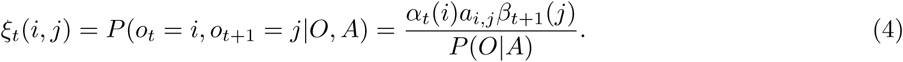

We then update the transition probabilities for each round according to (see Supplementary Information):

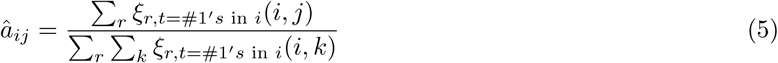

To assess convergence of the algorithm, we calculate the maximum change in any transition probability. For all of the cases in the results section we have used a convergence criterion of 0.001, so that if no transition probability changes by more than an absolute value of 0.001 between one iteration and the next we terminate the algorithm.

### Bootstrap resampling

We used 100 bootstrap resamples of observed transitions for estimating uncertainty in the HBW algorithm, and report the standard deviation of summary statistics over the set of resamples.

### HyperTraPS

HyperTraPS was implemented following [22], using 2 *×* 10^3^ samples to estimate likelihoods and a reduced parameterisation mapping *L*^2^ values to the edges of the hypercubic transition network. Priors involved a uniform probability of any remaining feature being acquired at any time step. The results from HyperTraPS were obtained after preliminary investigation to find the optimal perturbation kernel for MCMC convergence. For the simple synthetic cases this was *σ* = 0.75; for the ovarian and TB cases it was *σ* = 0.25.

### Summary of inferred dynamics with ordering probabilities

To summarise HyperHMM outputs, 10^5^ random walkers are simulated on the maximum likelihood hypercube, and the ordering of trait acquisition for each walker is recorded. Where the bootstrap is used, this process is repeated for each resampled dataset, and the standard deviation of each ordering probability over the resampled set is recorded. To summarise HyperTraPS outputs, 10^3^ random walkers are simulated on each of 10^5^ sampled hypercubes from the posterior, and the corresponding sampled trait-ordering posteriors are recorded.

### Ancestral reconstruction for tuberculosis dataset

Here we followed the minimum evolution approach in Ref. [22], where the ancestor of two descendants was inferred to possess a trait iff both descendants also possess it. This approach assumes that the acquisition of traits is rare, so convergent acquisition is correspondingly rare; in cases where this assumption is not safe, our approach can readily be applied across an ensemble of possible phylogenies and the resulting inferred dynamics summarised accordingly.

### Implementation

The implementation of the code is currently in C++. Currently the most effective implementation of the HyperHMM algorithm works on a system of at least 20 traits on a normal laptop. The implementation is made using the Compressed Row Storage, CRS, format for representing sparse matrices. Code for inference and visualisation is freely available at https://github.com/StochasticBiology/hypercube-hmm.

## Acknowledgments

This work was supported by the Trond Mohn Foundation (project HyperEvol under Grant Agreement No TMS2021TMT09), through the Centre for Antimicrobial Resistance in Western Norway (CAMRIA), Grant Number: TMS2020TMT11. This project has received funding from the European Research Council (ERC) under the European Union’s Horizon 2020 research and innovation programme [Grant agreement No. 805046 (EvoConBiO)]. We are grateful to Kostas Giannakis for helpful discussions.

---

**Figure S1:**
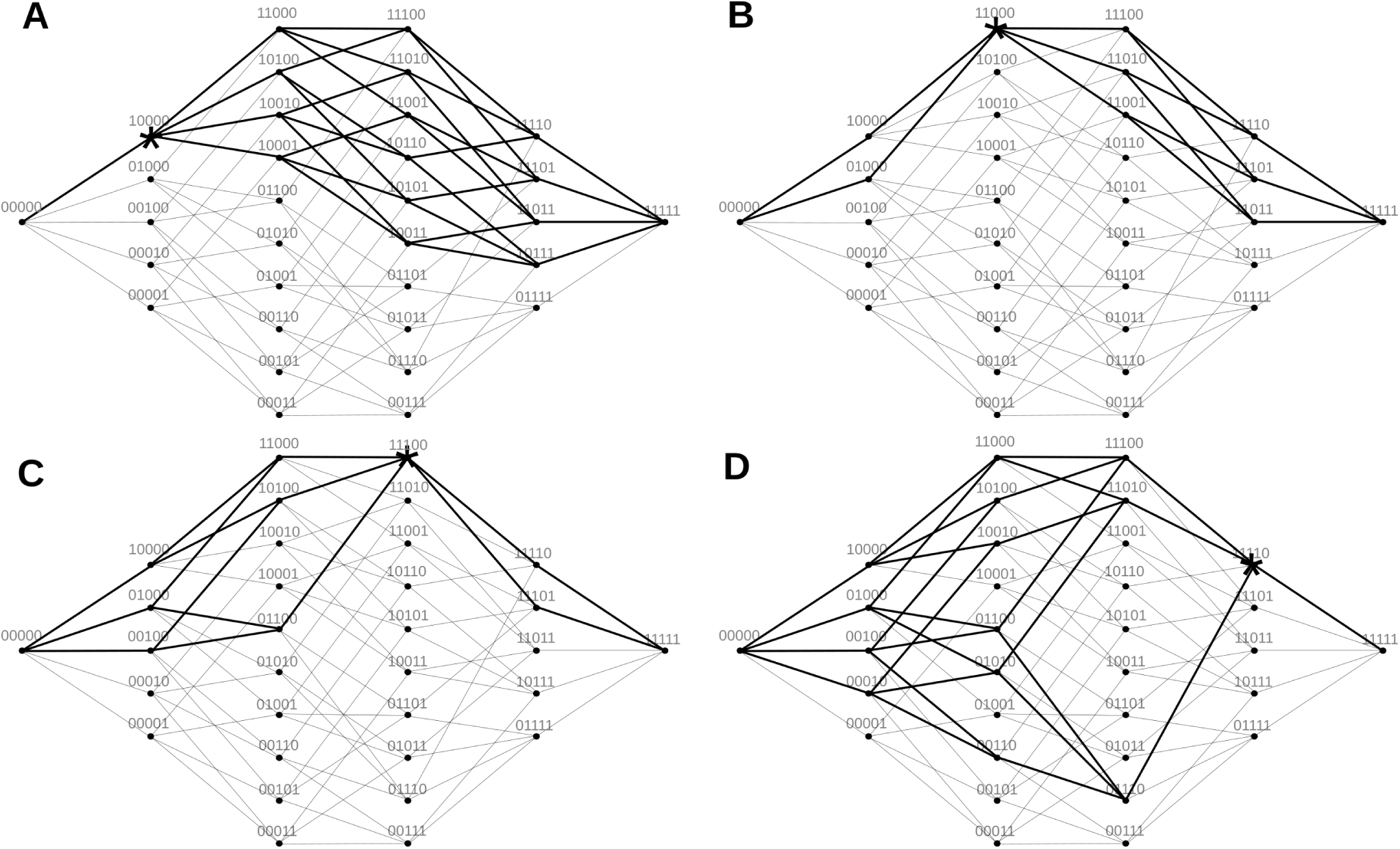
Illustration of hypercubic trajectories associated with an observation. Visualisation of all possible pathways given an observation on a 5D hypercube, where the black lines shows the possible pathways given the observations (marked with stars) (A) 10000, (B) 11000, (C) 11100, (D) 11110.

## A Supplementary Information

### A.1 Model setup

#### Definition A.1

(State space and observation sequences). The **state space** is a set consisting of all the possible states in which the system can exist. An **observation sequence** is a time ordered sequence of observed states from a given state space, denoted as *O*.

#### Definition A.2

(Transition probabilities and transition probability matrix [34]). A **transition probability** is the probability of going from one state to another in a stochastic process. Let *{X*_*t*_, *t* ∈ *T}* be a stochastic process with state space *S* = *s*_1_, *s*_2_, …, *s*_*N*_, then the corresponding transition probabilities are *P* (*X*_*n*_ = *s*_*j*_|*X*_*n−*1_ = *s*_*i*_) = *a*_*ij*_, each of which is the probability of going from state *s*_*i*_ to state *s*_*j*_. Note that 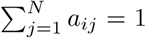, since we have to go to one of the states, and hence the probability of going from state *i* to any other state has to be 1.

The **transition probability matrix**, or just the transition matrix, is the matrix representation of all the transition probabilities:

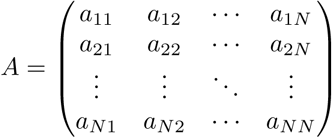

#### Definition A.3

(Emission probabilities and emission matrix [35]). Let *{X*_*t*_, *t* ∈ *T}* be a stochastic process with state space *S*, and *O* be the sequence of observations drawn from the set *{y*_1_, …, *y*_*m*_*}*. Then *b*_*i*_(*y*_*k*_) = *P* (*y*_*k*_|*X*_*n*_ = *s*_*i*_) denotes the **emission probabilities** (or observation likelihoods). This is the probability of generating observation *y*_*k*_ given that we are in state *s*_*i*_. Note that 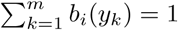.

The **emission matrix** is the matrix representation of all the emission probabilities:

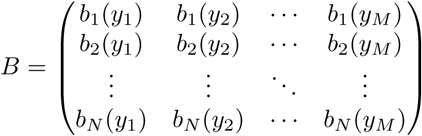

### A.2 The Baum-Welch algorithm

A hidden Markov model (HMM) is a model wherein a system evolves according to an unobservable Markov chain, but emits signals that are observable. A given HMM is described by the *A* and *B* matrices above. The Baum-Welch algorithm, developed in the late 1960’s and early 1970’s, is a variant of the expectation-maximisation algorithm that estimates transition- and emission probabilities in a hidden Markov model, with very diverse use across fields including speech recognition, computational biology, computer vision and econometric [36].

To use the Baum-Welch algorithm one need a finite number of states and a sequence, or multiple sequences, of observations. The goal of the Baum-Welch algorithm is to use this information to estimate the transition probabilities in the underlying Markov chain, and the emission probabilities connected to the Markov chain. One important feature of the Baum-Welch algorithm is that it is only guaranteed to find a local optimum, not necessarily the global optimum, as is the case with many learning algorithms [36]. This means that we are not guaranteed the best solution every time, and that the result might depend on the initialisation of the algorithm.

We proceed by reviewing the original Baum-Welch algorithm, before describing the adaptations to the multiple signals, hypercubic case. At the core of the algorithm are several sub-functions, which are called *α*-, *β*-, *ξ*-, and *γ*-function, also referred to as *α*-, *β*-, *ξ*-, and *γ*-probabilities [37]. We write *λ* = *A, B* for the set of parameters describing a particular HMM, and *q*_*t*_ for the state of the (unobservable) HMM at time *t*.

#### Definition A.4

(*α*-probability). The *α***-probability** is the probability of seeing a given set of observations up to time *t* and that we are in state *i* at time *t* given all of the hidden Markov model parameters *λ*:

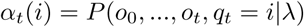

#### Definition A.5

(*β*-probability). Given a state *i* at time *t* and the model parameters *λ*, the *β***-probability** is the probability of observing all the observations starting from *o*_*t*+1_, the observation at time *t* + 1, and going to *o*_*T*_, the observation at *T* :

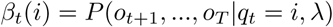

#### Definition A.6

(*ξ*-probability). Given an observation sequence *O* = *o*_0_, …, *o*_*T*_ and the model parameters, the *ξ***-probability** is the probability of being in state *i* at time *t* and state *j* at time *t* + 1:

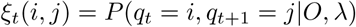

#### Definition A.7

(*γ*-probability). Given an observation sequence *O* = *o*_0_, …, *o*_*T*_ and the model parameters, the *γ***-probability** is the probability of being in state *i* at time *t*

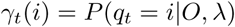

The *ξ*-function and *γ*-function is built up using the *α*-function and *β*-function, while the updating of transition and emission probabilities are done by using the *ξ*-function and *γ*-function.

The main idea of the Baum-Welch algorithm is to calculate the probability of seeing the observations when moving forward in time and when moving backward in time independently using the initial estimate of *A* and *B*. Then these calculations can be used to find a value for being at any given state at a given time. We can then normalize these values to find the new estimate of *A* and *B*. Iterating this process repeatedly until convergence is the Baum-Welch algorithm, given in Algorithm 2, with each step explained in more detail in Section A.4.

### A.3 Multiple Sequence Baum-Welch Algorithm

The original Baum-Welch algorithm takes one observation sequence, *O* = *o*_0_, …, *o*_*T*_. In hypercubic inference we typically have a dataset involving multiple independent evolutionary or progressive instances of a system, and are therefore interested in estimating parameters that best describe the dynamics of this ensemble of instances. We therefore require a multiple-sequence adaptation of the Baum-Welch algorithm.

#### Algorithm 2

The Baum-Welch algorithm [35]. Takes a set of *T* + 1 observations *o*_0_, …, *o*_*T*_, estimates transition matrix *A* and emission matrix *B*.

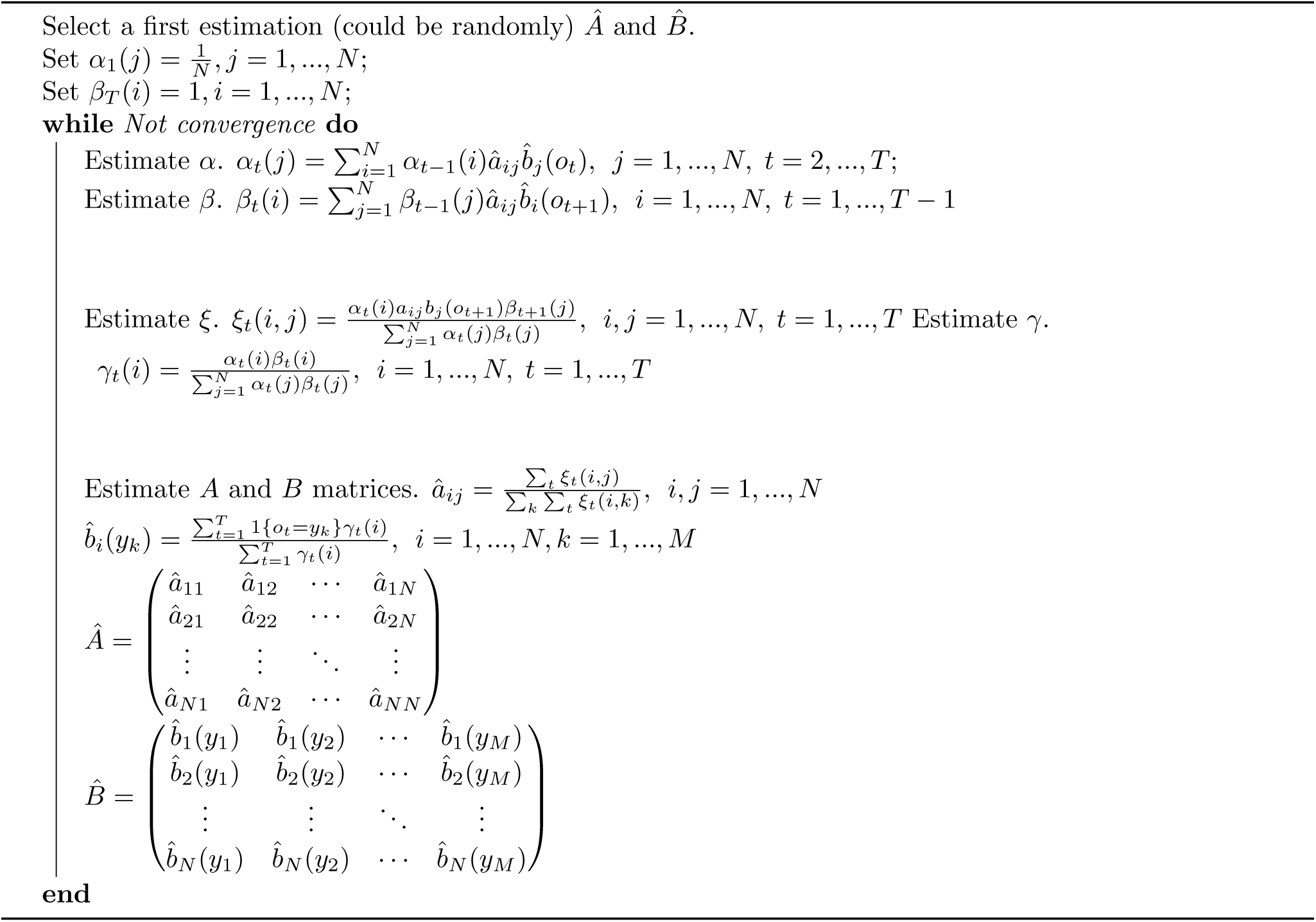

In the single-sequence Baum-Welch we update the transition probabilities using the following formula:

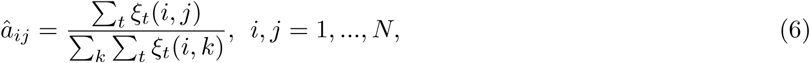

estimating the probability, summed over all times, of being in *j* the timestep after being in *i*, normalised by the total probability of being in any state *k* the timestep after being in *i*.

Let us use the following notation, *O*_*r*_ = *o*_*r*,0_, …, *o*_*r,T*_, to denote the *r*’th observation sequence, and *ξ*_*rt*_ is the *ξ*-probability calculated using *O*_*r*_ at time *t*. Note that all *O*_*r*_ is independent of each other. Then we can update the transition probabilities like this instead:

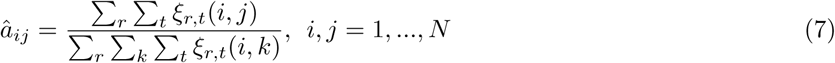

Which is the same as before, except that we now sum over all the independent observation sequences as well.

### A.4 Hypercubic Multiple Sequence Baum-Welch Algorithm

Our main objective is to infer transition probabilities over a directed hypercube given some data. The weight of the edges on the hypercube is the transition probabilities from node *i* to node *j*. The Markov condition, that the probability of going from one state to another is only dependent on the previous state, holds in this case, so we can fairly assume that a walk on this hypercube is a Markov chain.

The state space in this case is the set of all nodes on the hypercube, meaning if we have *L* traits there are 2^*L*^ possible states to be in. With a complete transition graph this would imply an extremely large transition matrix for high *L*, with 2^*L*^ *·* 2^*L*^ values. However, since we are assuming a hypercubic structure – that is, that no entity can remove a trait once they have it, and that we can only get one trait at a time – this transition matrix is sparser. The number of allowed transitions are the number of edges on the hypercube, 2^*L−*1^*L*, which means we will have 2^2*L*^ *−* 2^*L−*1^*L* zero entries in the matrix.

The observation data we are considering will typically involve transitions between two states. This will either be the transition from the 0^*L*^ initial state to a given observation (in the case of cross-sectional data) or the transition from some precursor state to some subsequent state (in the case of longitudinal or phylogenetic data).

We begin by picturing each independent observed transition as a subset of a longer trajectory from the node of all zeroes to the node of all ones. As steps through our hypercubic state space involve the acquisition of one trait at a time, each such long trajectory will involve *L* steps. We use the wildcard character ‘?’ to represent all unspecified states in a given trajectory. For example, a cross-sectional observation of 1001 would give the trajectory 0000 →? → 1001 →? → 1111, and a longitudinal observation 1100 → 1101 would give 0000 →? → 1100 → 1101 → 1111.

Writing the observation set in this way gives us some advantages. First, notice that *o*_0_ always equals all 0’s and *o*_*T*_ always equals all 1’s, and that every observation sequence, once we have added the unknown states, are of the same length. The question mark will from now on represent all possible states to be in at that time given the observation sequence.

Let us start by looking at how we can calculate the *α*-probabilities. In the normal Baum-Welch algorithm the *α*-probabilities is defined as follow, *α*_*t*_(*i*) = *P* (*o*_0_, …, *o*_*t*_, *q*_*t*_ = *i* |*λ*). Since *o*_*t*_ is just a given state at time *t*, this gives *α*_*t*_(*i*) = *P* (*o*_0_, …, *o*_*t*_ = *i* |*λ*). At *t* = 0 we have *α*_0_(*i*) = *P* (*o*_0_ = *i* |*λ*) = 1 for *i* = 0^*L*^ (the initial state), and 0 for all other states.

For *t >* 0, we know that *α*_*t*_(*i*) = *P* (*o*_0_, …, *o*_*t*_ = *i*). Marginalising on the final step, it follows that *P* (*o*_0_, …, *o*_*t*_ = *i*) = Σ_*j*_ *P* (*o*_0_, …, *o*_*t−*1_ = *j*) *· P* (*j* → *i*). Here *P* (*o*_0_, …, *o*_*t−*1_ = *j*) = *α*_*t−*1_(*j*) and *P* (*j* → *i*) = *a*_*j,i*_. This gives the following recursive formula for calculating *α*_*t*_(*i*) when *t >* 0:

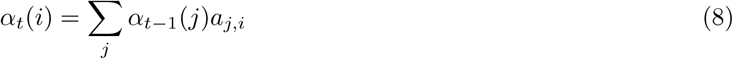

In an unrestricted case, calculating and summing this value for every state would become intractable for large *L*. The hypercubic structure of the problem dramatically simplifies the calculation. Let us first look at the sum over all states *j*. Here, *a*_*j,i*_ = 0 for every state *j* where it is not possible to move from *j* to *i*. This means that we only need to sum over all states where it is possible to move from *j* to *i*. Let us define this set as follows:

#### Definition A.8.

*j* ∈ *D*_*i*_ if, and only if, *a*_*j,i*_ *>* 0

If we know the state at time *t*, we only need to calculate *α*_*t*_(*i*) for that given state and set all other *α*_*t*_ values to 0. However, if we do not know the state at time *t*, we need to calculate *α*_*t*_(*i*) for every *i* accessible from *j* where *α*_*t−*1_(*j*) *>* 0. This set of accessible *i* states will be defined as *F*_*t*_ as follows:

#### Definition A.9.

*i* ∈ *F*_*t*_ if and only if there exists a state *j* such that *a*_*j,i*_ *>* 0 and *α*_*t−*1_(*j*) *>* 0.

This gives us the following expression for *α*_*t*_(*i*).

#### Definition A.10.

[*α*-probabilities]

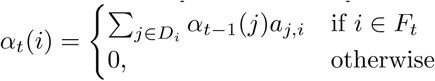

This calculation only applies to the case where *o*_*t*_ is unknown. If *o*_*t*_, is known we would know that every *α*_*t*_(*i*) = 0 as long as *i ≠ o*_*t*_. The calculation for *α*_*t*_(*o*_*t*_) is however the same as the upper equation in Def. A.10.

The *β*-probabilities are calculated in a similar way. We start by looking at the most simple case, when *t* = *T*. Since we do not have any observations after time *T*, we will define *β*_*T*_ (*i*) to be 1 for every *i* ∈ *S*.

For *t < T* we have the following expression in the normal Baum-Welch algorithm: *β*_*t*_(*i*) = *P* (*o*_*t*+1_, …, *o*_*T*_ *o*_*t*_ = *i, λ*). This equation can be rewritten such that we gain a recursive formula, like in Def. A.10.

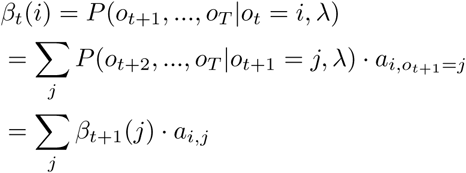

Like with the *α*-probabilities we do not need to calculate everything all the time, but the general rule for calculating the *β*-probabilities is:

#### Definition A.11.

Let *B*_*t*_ denote the set of all possible states to be in at time *t*. The set *B*_*t*_ hence contains all binary strings which has *t* 1’s.

#### Definition A.12.

[*β*-probabilities] 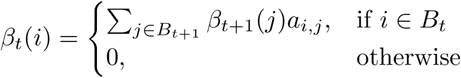, for *T > t*

For the *ξ*-probabilities we have to do something slightly different. The most simple case is when both *o*_*t*_ and *o*_*t*+1_ are known. When this is the case, we have full information and know that *ξ*_*t*_(*i, j*) = 1 when *i* = *o*_*t*_ and *j* = *o*_*t*+1_. For every other value of *i* and *j, ξ*_*t*_(*i, j*) = 0.

In general, *ξ*_*t*_(*i, j*) can be calculated using the *α*- and *β*-probabilities. We know the probability of seeing everything up to time *t*, and being in state *i* at time *t* (*α*-probabilities), we know the probability of going from *i* to *j* (*a*_*i,j*_), and we know the probability of seeing everything after time *t* given state *j* (*β*-probabilities). Then *ξ*_*t*_(*i, j*) will just be all these probabilities multiplied together.

As with the *α*- and *β*-probabilities we can use the known observation sequence to cut down the number of needed calculations. When *o*_*t*_ is known and *o*_*t*+1_ is unknown, we only need to calculate for every *j* which is possible to reach from *i*. If the opposite is true, that we know *o*_*t*+1_ but not *o*_*t*_, we only need to calculate for every *i* where it is possible to go to *j*.

The last possible case is when none of the states are known. In this case we need to calculate *ξ*_*t*_(*i, j*) for every state *i* which is possible to be in at time *t* and every state it is possible to go to from that state.

#### Definition A.13

(*ξ*-probabilities). For the *ξ*-probabilities we will have these four cases: If both *o*_*t*_ and *o*_*t*+1_ are known:

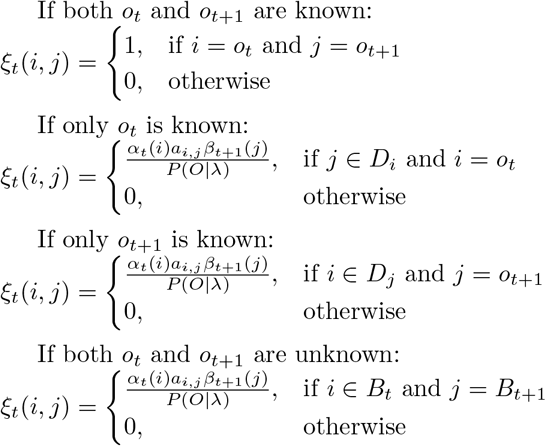

**Figure S2:**
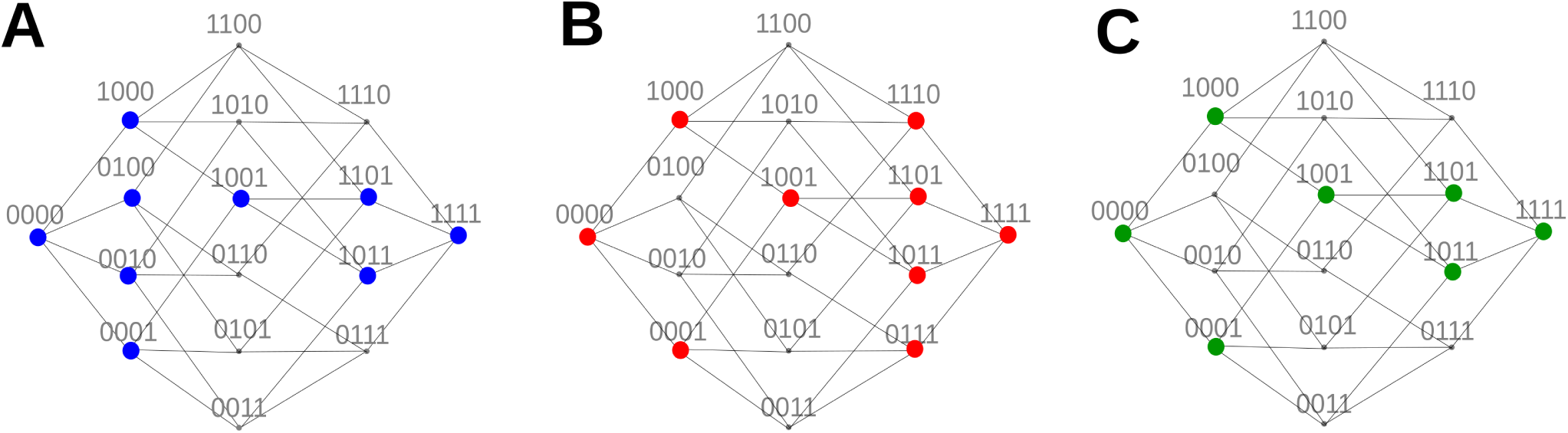
Example nodes considered in the HBW calculation. Given the observation 1010, (A) shows nodes that will have *α*_*t*_(*i*) *>* 0 for any time *t*; (B) shows nodes that will have *β*_*t*_(*i*) *>* 0 for any time *t*; (C) shows nodes that are present in both (A) and (B). All the edges connecting two nodes here will be given some weight when uploading the transition probabilities.

Now to the updating of the transition probabilities. We intialise the algorithm with a uniform transition matrix, meaning each value in the matrix is 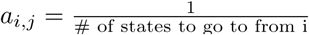, if it is possible to go from *i* to *j*, and 0 otherwise. We then need to calculate the *ξ*-probabilities for all of the given observation sequences. When we have the *ξ*-probabilities we can update our transition matrix. This is done the following way:

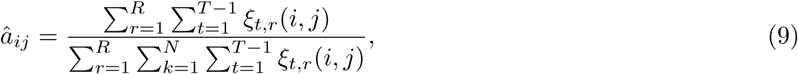

where *ξ*_*t,r*_(*i, j*) is *ξ*_*t*_(*i, j*) for observation sequence *r*.

To illustrate the subsets of space involved in a calculation, Figure S2 shows all the nodes which will be given a value *>* 0 for (a) the *α*-probabilities, (b) the *β*-probabilities, and (c) the *ξ*-probabilities for the observation 1010.

## B Hypercubic Baum-Welch example calculation

To illustrate the algorithm, we now consider an explicit example. Assume we are given 10100 as an observation in a cross-sectional context. That is interpreted as the sequence 00000 → ? → 10100 → ? → ? → 11111. We begin with a uniform transition matrix, and proceed to calculate the *α*-, *β*-, and *ξ*-probabilities. Calculations are described one timepoint at a time in the text, and illustrated in Fig. S3.

### B.1 *α* calculations

We first need to calculate the forward probabilities, defined as follows:

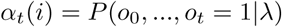

For *t* = 0 we have *α*_0_(00000) = 1 and *α*_0_(*i*) = 0, for all *i* except 00000. This will always be the case independent of the observation.

For *t* = 1 we will have *α*_1_(*i*) = *P* (*o*_0_ = 00000, *o*_1_ = *i*) = *P* (00000 → *i*). This will only have a value for *i* ∈ *{*10000, 01000, 00100, 00010, 00001*}* since we are assuming that we only acquire one trait at a time. For the case where *i* ∈ *{*10000, 01000, 00100, 00010, 00001*}* we have *α*_1_(*i*) = *P* (00000 → *i*) = *a*_00000,_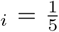, and for everything else we have *α*_1_(*j*) = 0.

For *t* = 2 we have *α*_2_(10100) = *P* (*o*_0_ = 00000, *o*_1_, *o*_2_ = 10100) = Σ_*j*_ *P* (*o*_0_ = 00000, *o*_1_ = *j*)*P* (*j* → 10100) =Σ_*j*_ *α*_1_(*j*) *· a*_*j*,10100_. Here only *j* ∈ *{*10000, 00100*}* give a non-zero value for both *α*_1_(*j*) and *a*_*j*,10100_. Hence, we end up with 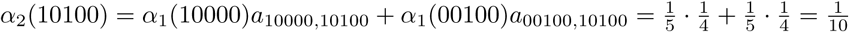.

For *t* = 3 we have *α*_3_(*i*) = *P* (*o*_0_ = 00000, *o*_1_, *o*_2_ = 10100, *o*_3_ = *i*) = *P* (*o*_0_ = 00000, *o*_1_, *o*_2_ = 10100) *· a*_10100,*i*_ = *α*_2_(10100) *· a*_10100,*i*_. Here only the cases where it is possible to go from 10100 will have non-zero values. Hence, for *i* ∈ *{*11100, 10110, 10101*}* we will have 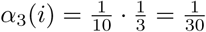.

For *t* = 4 we have *α*_4_(*i*) = *P* (*o*_0_ = 00000, *o*_1_, *o*_2_ = 10100, *o*_3_, *o*_4_ = *i*) = Σ_*j*_ *P* (*o*_0_ = 00000, *o*_1_, *o*_2_ = 10100, *o*_3_ = *j*) *· a*_*j,i*_ = Σ_*j*_ *α*_3_(*j*) *· a*_*j,i*_. Here again we will only have a few possible non-zero values. Since *α*_3_(*j*) ≠ 0 only for *j* ∈ *{*11100, 10110, 10101*}* only the places we can go from these three will have non-zero values for *α*_4_. Hence, if *i* ∈ *{*11110, 11101, 10111*}* then 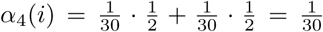. This is happening since for all *j* ∈ *{*11100, 10110, 10101*}* there is only two possible ways to go.

For *t* = 5 we have 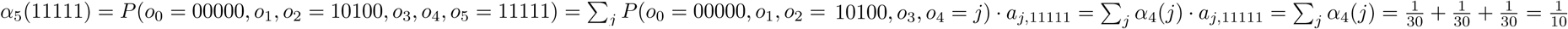. For *t* = 5 everything else will be 0.

### B.2 *β* calculations

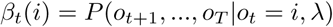

For *t* = 5 we have *β*_5_(*i*) = 1 for every *i*.

For *t* = 4 we have *β*_4_(*i*) = *P* (*o*_5_|*o*_4_ = *i*) = *P* (11111|*o*_4_ = *i*) = *a*_*i*,11111_. If *i* ∈ *{*11110, 11101, 11011, 10111, 01111*}* we will have *β*_4_(*i*) = 1. For everything else *β*_4_(*i*) = 0.

For *t* = 3 we have *β*_3_(*i*) = *P* (*o*_4_, *o*_5_ = 11111|*o*_3_ = *i*) = Σ_*j*_ *P* (*o*_5_ = 11111|*o*_4_ = *j*) *· a*_*i,j*_ *Σ*_*j*_ *β*_4_(*j*) *· a*_*i,j*_. From *t* = 4 we have that *β*_4_(*j*) = 1 for *j* ∈ {11110, 11101, 11011, 10111, 01111} and 0 otherwise. Hence we only need to calculate the values for every state where it is possible to reach one of these five values. Since these five values are all the possible states to be in at time *t* = 4 and the transition matrix is uniform 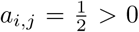 for every state that is possible to be in at time *t* = 3. We then end up with 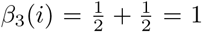 for all *i* which is possible to be in at time *t* = 3.

For *t* = 2 we have *β*_2_(10100) = *P* (*o*_3_, *o*_4_, *o*_5_ = 11111|*o*_2_ = 10100) = Σ_*j*_ *P* (*o*_4_, *o*_5_ = 11111|*o*_3_ = *j*) *· a*_10100,*j*_ = Σ _*j*_ *β*_3_(*j*) *· a*_10100,*j*_. Here we can see that this will only have a non-zero value for states it is possible to go from 10100, and we know from time *t* = 3 that all non-zero values of *β*_3_ is 1. Hence, we get 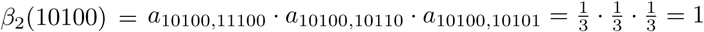.

For *t* = 1 we have *β*_1_(*i*) = *P* (*o*_2_ = 10100, *o*_3_, *o*_4_, *o*_5_ = 11111|*o*_1_ = *i*) = *P* (*o*_3_, *o*_4_, *o*_5_ = 11111|*o*_2_ = 10100) *· a*_*i*,10100_ = *β*_2_(10100) *· a*_*i*,10100_ = *a*_*i*,10100_. This will only have a non-zero value for states where it is possible to go to 10100. Hence, if *i* ∈ *{*10000, 00100*}* we have 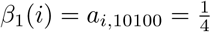.

For *t* = 0 we have *β*_0_(0 0000) = *P* (*o*_1_, *o*_2_ = 10100, *o*_3_, *o*_4_, *o*_5_ = 11111|*o*_0_ = 00000) = Σ _*j*_ *P* (*o*_2_ = 10100, *o*_3_, *o*_4_, *o*_5_ = 11111|*o*_1_ = *j*) *·a*_00000,*j*_ = Σ_*j*_ *β*_1_(*j*)*·a*_00000,*j*_. We know from time *t* = 1 that *β*_1_(*j*) *>* 0 only for *j* ∈ *{*10000, 00100*}*. Hence, we get 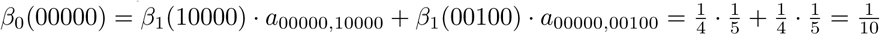.

### B.3 *ξ* calculations

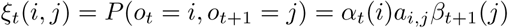

For *t* = 0 we have *ξ*_0_(00000, *j*) = *P* (*o*_0_ = 00000, *o*_1_ = *j*). This will only be non-zero for values where it is possible to go to from 00000. We also see that from the *β*-calculations that only the states 10000 and 00100 have non-zero values for *β*_1_ so we only need to look at these two states. Again, since our transition matrix is uniform so every calculation will be similar. So, if *j* ∈ *{*10000, 00100*}* we have 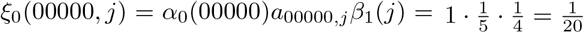. For every other combination of *ξ*_0_(*i, j*) = 0.

**Figure S3:**
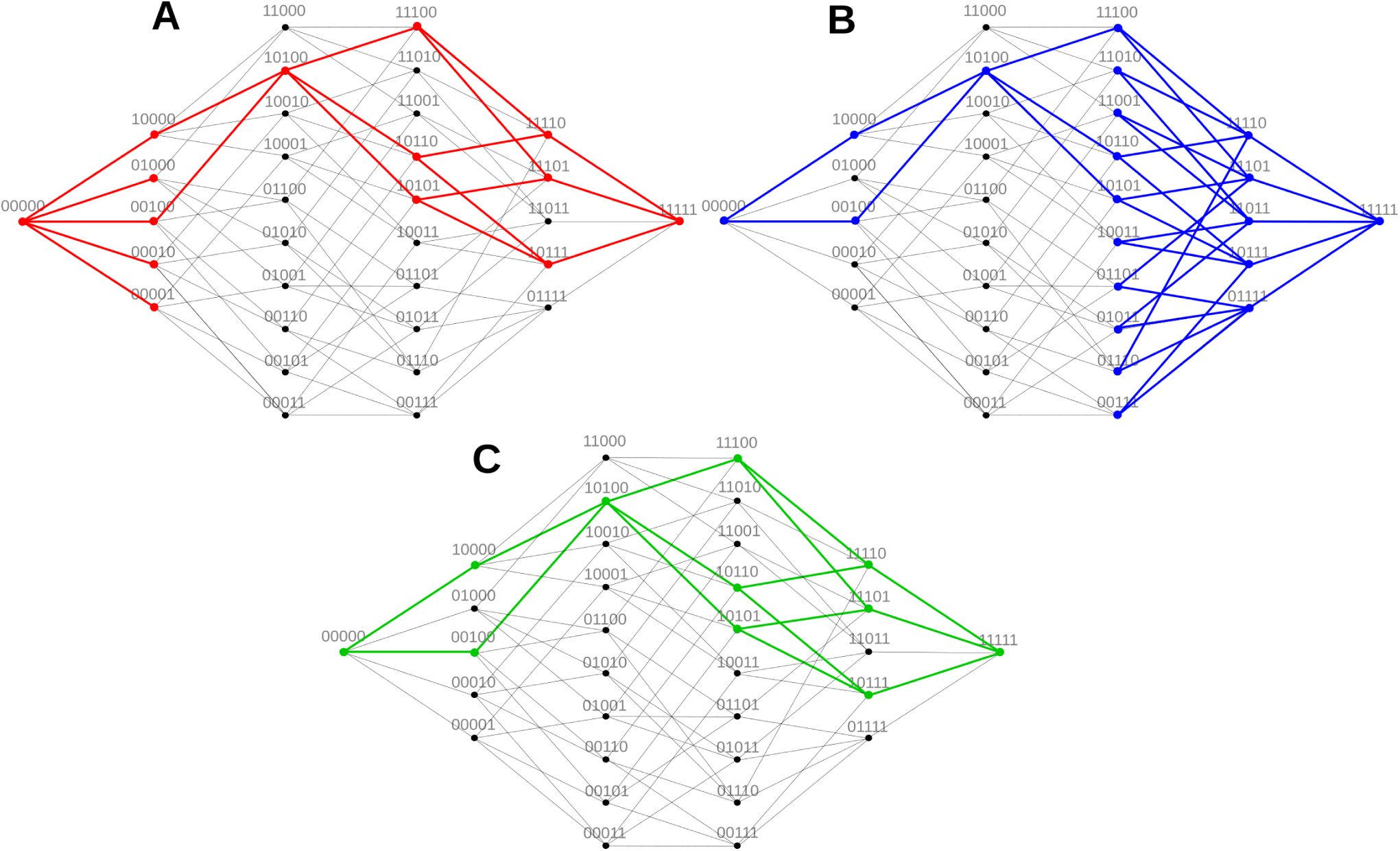
Example edges considered in the HBW calculation. Given the observation 10100, (A) shows all the nodes that will have *α*_*t*_(*i*) *>* 0 for any time *t*; (C) shows all the nodes that will have *β*_*t*_(*i*) *>* 0 for any time *t*; (C) shows all the nodes that are included in both (A) and (B). All the edges connecting two nodes here will be given some weight when uploading the transition probabilities.

For *t* = 1 we have *ξ*_1_(*i*, 10100) = *P* (*o*_1_ = *i, o*_2_ = 10100) = *α*_1_(*i*)*a*_*i*,10100_*β*_2_(10100). From the *α*-calculations we know that 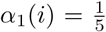 for *i* ∈ *{*10000, 01000, 00100, 00010, 00001*}*, but here we also see that only 10000 and 00100 will give a non-zero value for *a*_*i*,10100_. Hence, we end up with 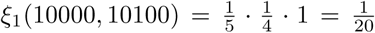 and 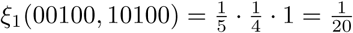.

For *t* = 2 we have *ξ*_2_(10100, *j*) = *P* (*o*_2_ = 10100, *o*_3_ = *j*) = *α*_2_(10100)*a*_10100,*j*_*β*_3_(*j*). Since all states which is possible to be in at time *t* = 3 have a value for *β*_3_ we only need to consider the allowed transitions from 10100. Which gives us, for *j* ∈ *{*11100, 10110, 10101*}* we have 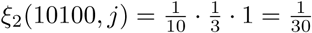.

For *t* = 3 we have *ξ*_3_(*i, j*) = *P* (*o*_3_ = *i, o*_4_ = *j*) = *α*_3_(*i*)*a*_*i,j*_*β*_4_(*j*). Again, from the *α*-calculations we have that *α*_3_(*i*) *>* 0 for *i* ∈ *{*11100, 10110, 10101*}*. For *j* we will have non-zero values when it is possible to go from on of the states *i*. So, if *i* ∈ *{*11100, 10110, 10101*}* and *j* ∈ *{*11110, 10111, 11101*}* we have 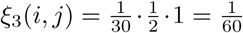.

For *t* = 4 we have *ξ*_4_(*i*, 11111) = *P* (*o*_4_ = *i, o*_5_ = 11111) = *α*_4_(*i*)*a*_*i*,11111_*β*_5_(11111). Here the calculations are restricted by the non-zero values of *α*_4_(*i*). For *i* ∈ *{*11110, 10111, 11101*}* we have 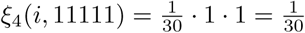.

## C Library credits

The inference code uses the C++ Armadillo library [38]. Bubble plot visualisations use Python libraries matplotlib [39], pandas [40], seaborn [41], and numpy [42]. Other visualisations use R libraries stringr [43], ggplot2 [44], ggrepel [45], gridExtra [46], and igraph [47].

